# Dissecting earthworm diversity in tropical rainforests

**DOI:** 10.1101/2024.09.13.612984

**Authors:** Arnaud Goulpeau, Mickaël Hedde, Pierre Ganault, Emmanuel Lapied, Marie-Eugénie Maggia, Eric Marcon, Thibaud Decaëns

## Abstract

Tropical rainforests are among the most emblematic ecosystems in terms of biodiversity. However, our understanding of the structure of tropical biodiversity is still incomplete, particularly for certain groups of soil organisms such as earthworms, whose importance for ecosystem functioning is widely recognised. This study aims at determining the relative contribution of alpha and beta components to earthworm regional diversity at a hierarchy of nested spatial scales in natural ecosystems of French Guiana. For this, we performed a hierarchical diversity partitioning of a large dataset on earthworm communities, in which DNA barcode-based operational taxonomic units (OTUs) were used as species surrogates. Observed regional diversity comprised 256 OTUs. We found that alpha diversity was lower than predicted by chance, regardless of the scale considered. Community-scale alpha diversity was on average 7 OTUs. Beta diversity among remote landscapes was higher than expected by chance, explaining as much as 87% of regional diversity. This points to regional mechanisms as the main driver of species diversity distribution in this group of organisms with low dispersal capacity. At more local scales, multiplicative beta diversity was higher than expected by chance between habitats, while it was lower than expected by chance between communities in the same habitat. This highlights the local effect of environmental filters on the species composition of communities. The calculation of a Chao 2 index predicts that as much as 1,700 species could be present in French Guiana, which represents a spectacular increase compared with available checklists, and calls into question the commonly accepted estimates of global number of earthworm species.

## Introduction

Tropical rainforests are among the most emblematic ecosystems in terms of biodiversity. They make up fifteen of the World’s twenty-five terrestrial biodiversity hotspots, i.e. regions with an exceptional concentration of endemic species that are also suffering exceptional habitat loss (Myers et al. 2000). One hectare of tropical rainforest generally contains at least 100 tree species, with extreme values of over 500 (Valencia et al. 1994), and it is widely accepted that these ecosystems host more than 50% of the Earth’s species on just 7% of the Earth’s land surface (Wilson 1988). To date, understanding the mechanisms underlying this diversity remains a major scientific challenge, and a multitude of hypotheses have been put forward to explain it (Hill and Hill 2001, Willig et al. 2003, Dyer et al. 2007). These hypotheses differ in the time step they considered (i.e. ecological, biogeographical or evolutionary), in the consideration they give to non-deterministic processes (i.e. neutralist theory of biodiversity), and in the component of diversity they emphasise (i.e. alpha, beta or gamma component sensu Whittaker 1972).

It is commonly accepted that these diversification mechanisms, whatever the one involved, lead to a structuring of species diversity in which most of the regional diversity is explained by the beta component (Erwin 1982, Condit et al. 2002, Willig et al. 2003, Beck et al. 2012). However, this importance is also likely to vary geographically (Condit et al. 2002, Dahl et al. 2009, Willig and Presley 2013), and the generality of this pattern has also been challenged by studies on understory phytophagous insects (Novotny et al. 2007, Hulcr et al. 2008). These apparent contradictions may reflect different mechanisms involved in structuring the diversity of different groups of organisms, but they may also result from differences in observation scales and sampling designs between studies and to undersampling bias that can lead to an overestimation of the beta component (Condit et al. 2002, Novotny et al. 2007, Beck et al. 2012). Another obstacle to our understanding of the structuring of tropical biodiversity lies in the considerable imbalance of knowledge in favour of a small number of well-known organisms, essentially plants and a few animal taxa, which at best represent only a tiny fraction of tropical biodiversity. To date, these various limitations still prevent us from having a consolidated view of the relative importance of the alpha and beta components in explaining the peak in biodiversity observed in tropical rainforests.

In this study, we seek to understand the spatial structuring of earthworm diversity, a group of soil animals considered essential for the functioning of terrestrial ecosystems but paradoxically suffering from a considerable knowledge deficit, especially in the tropics where most of the species richness concentrates (Decaëns et al. 2006, Decaëns 2010, Phillips et al. 2019). In Amazonia, for example, it has been suggested that several hundred species, most of which have yet to be discovered, could exist in an area as small as French Guiana (Decaëns et al. 2016, 2024, Maggia et al. 2021). This high diversity is characterised by a particularly high beta component, both at the scale of Amazonia as a whole (Lavelle and Lapied 2003) and at a more regional scale (Maggia et al. 2021, Conrado et al. 2023). This pattern has been interpreted as the result of the low dispersal capacity of these organisms, which favours allopatric speciation mechanisms and prevents exchanges of species across biogeographic barriers that would be insignificant for other more mobile organisms (Bouché 1972, Lavelle and Lapied 2003).

We explored a substantial dataset accumulated over several expeditions to tropical rainforests of French Guiana (Goulpeau et al. 2024). We partitioned regional earthworm diversity to figure out the relative importance of its alpha and beta components through a hierarchy of nested spatial scales. We then used our results to propose an estimate of the number of earthworm species existing at a regional scale. Based on current knowledge of the diversity of Amazonian earthworms, we hypothesised: 1) that regional diversity is essentially explained by large-scale compositional dissimilarity, while alpha and local-scale beta components are of comparatively minor importance, and 2) that the studied region hosts an unsuspected number of earthworm species. This last hypothesis, if confirmed, would likely call into question the current consensus on the number of species that exist on a global scale.

## Materials and methods

### Study sites

Earthworm communities were sampled in French Guiana, an area of 83,846 km^2^ covered by over 95% of tropical rainforests, which is located in the eastern Guiana precambrian shield or the Amazonian basin, between the Oyapock and Maroni rivers. The climate is humid tropical, with an average annual temperature of around 26 °C and average annual rainfall ranging from 2,000 to 4,000 mm (Barret 2001). The rainy season extends from December to June, interrupted by a short dry period (typically two weeks) in February or March. The region is mainly lowlands, criss-crossed by a dense river network and interrupted by regions of isolated hills and inselbergs that can reach elevations of around 800 m (Barret 2001). The soils are mainly Ferralsols in well-drained areas, sometimes superficially hardened by laterites, Lithosols on granitic outcrops (inselbergs and other landforms), Acrisols in lowland areas, and Podzols in sandy coastal regions (Ferry et al. 2003, FAO 2015).

The sampling design of this study has been detailed in Goulpeau et al. (2024). Ten localities were sampled during the rainy season between 2011 and 2019 (Fig. 1; SI Table S1). In each locality, sampling was carried out in one-hectare plots, spaced at least 500 m apart, and representing the main habitat types of the local landscape (SI Table S2): plateau forests (dense rainforest on deep, well-drained soils, sometimes with superficial laterites), slope forests (rainforest on slopes and deep soils), lowland forests (periodically flooded rainforest on hydromorphic soils) and inselbergs habitats such as rocky savannas (herbaceous formations dominated by terrestrial bromeliads), transitional forests and hilltop forests (low canopy rainforest on shallow soils). In most cases, habitats were replicated at least three times in each locality, giving seven to 25 one-hectare plots per locality and a total of 125 plots.

**Figure 1.**
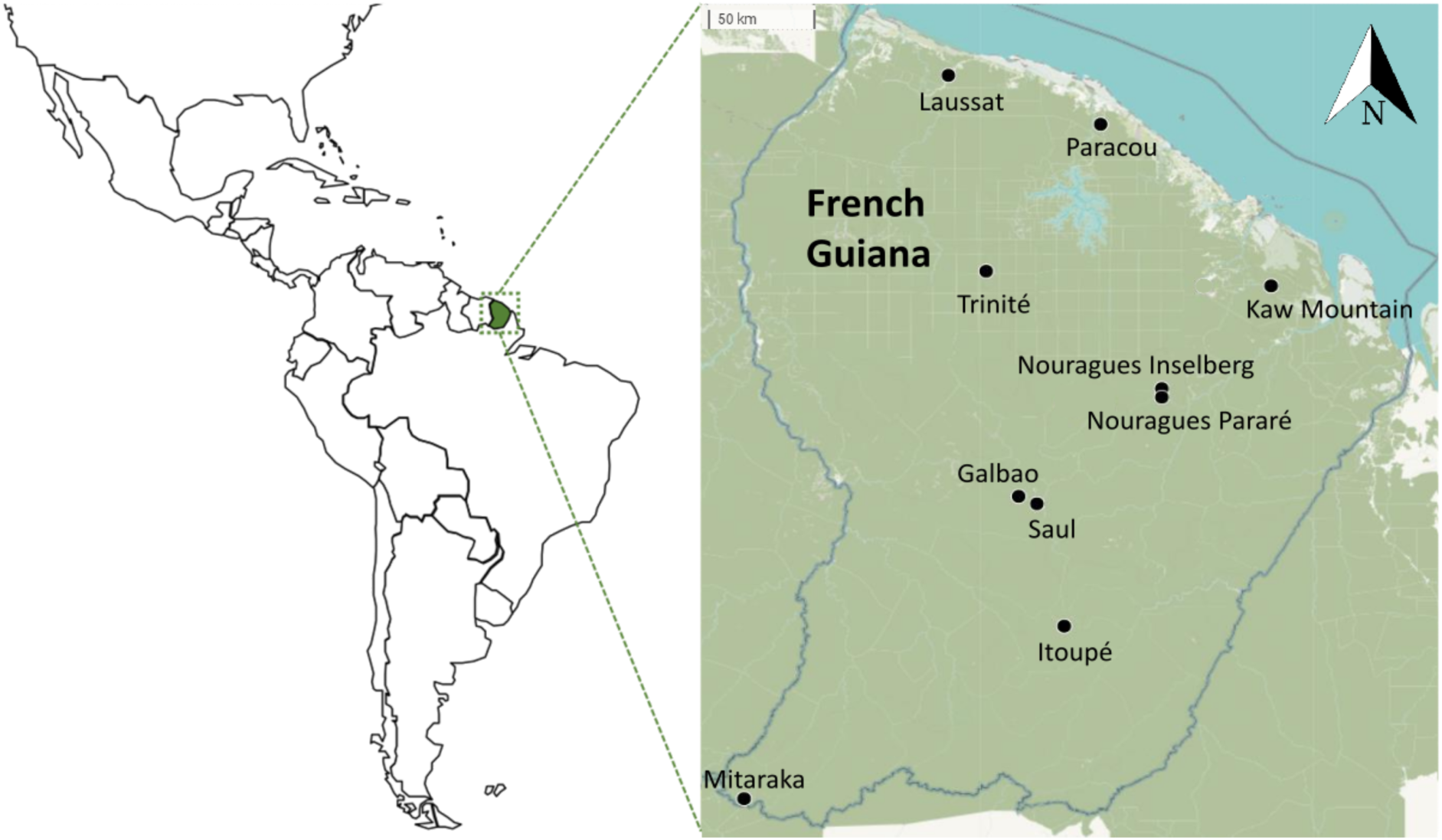
Location of the ten sampling localities in French Guiana.

### Earthworm sampling

On each one-hectare plot, earthworms were sampled using a combination of three methods (Decaëns et al. 2016, Maggia et al. 2021, Goulpeau et al. 2024): 1) three blocks of soil, 25 × 25-cm^2^ area and 20-cm deep, arranged on a central triangle of 20-m sides, were sorted manually on a plastic sheet; 2) for the larger soil-dwelling species, an area of 1 m^2^ was excavated to a minimum depth of 40 cm, selecting where possible an area showing traces of large earthworms on the soil surface (i.e. large casts); 3) finally, visual sampling was carried out on the one-hectare plot, looking for earthworms in all available and accessible microhabitats (i.e. decomposing trunks, litter accumulations, streamside sediments, epiphytes, etc.) for a fixed period of three hours.persons (e.g. one hour for three people). All life stages (i.e. adults, juveniles and cocoons) were collected, and specimens collected using each method and in each type of microhabitat were fixed separately in 100% ethanol. The ethanol was changed in each tube after a period of 24 h to ensure good fixation.

### DNA barcoding and OTU delimitation

The complete procedure for DNA sequencing and OUT delimitations has been described in Goulpeau et al. (2022). Briefly, for each sample (i.e. individuals collected with a given method in a given microhabitat and plot), earthworms were first sorted according to external morphological features (characteristics of the clitellum when adult, size and pigmentation), and for each morpho-group obtained, up to 5 individuals per sample were kept for molecular analysis. For the majority of samples (77.2%), DNA extraction, PCR reactions and sequencing of the barcode region of the Cytochrome C Oxidase I gene (COI) were carried out at the Centre for Biodiversity Genomics (Guelph, Canada) using standard protocols of the International Barcode of Life. A cocktail combining the M13 tail primer pairs LCO1490/HCO2198 (Folmer et al. 1994) and LepF1/LepR1 (Hebert et al. 2004) was used. Samples that failed after this first step were amplified using the internal primers MLepR1/MLepF1 and LCO/HCO pairs (Hajibabaei et al. 2006). The remaining sequences (22.8%) were obtained from the Laboratoire d’Ecologie Alpine (Grenoble, France). The LCO1490/HCO2198 primer pair (Folmer et al. 1994) was used with a label on the 5’ side to allow sample identification during bioinformatics analysis of the results, each sample being characterised by its labels on the forward and reverse primers. PCR products were sequenced on MiSeq (paired-read sequencing, 2 x 250 bp), and bioinformatic analysis of the sequences was performed using OBITools (Boyer et al. 2016; www.grenoble.prabi.fr/trac/OBITools). Approximately 220 bp were obtained for each end of the DNA barcode. All sequences are available in the BOLD dataset “Earthworms from French Guiana – 2023 update” (DS-EWFG2023; doi.org/10.5883/DS-EWFG2023).

Sequences were grouped into molecular operational taxonomic units (OTUs) using the Assemble Species by Automatic Partitioning (ASAP) method, which is the most suitable both for our study model and for the size of the dataset (Goulpeau et al. 2022). ASAP is based on an algorithm using only pairwise genetic distances in order to reduce the computation time for phylogenetic reconstruction (Puillandre et al. 2021). To do this, the algorithm ranks the dissimilarity matrix values in ascending order, with each value in turn becoming the limit value between intra– and interspecific divergence levels. It then proposes a series of different sequence groupings, each of which is assigned a score based on two criteria: the probability that a group of sequences represents a species (p-value) and the width of the sequence gap between the current state of the group and its state before the grouping (W). The p-values and W-values are then ranked in ascending and descending order respectively. For each delimitation, the average of the ranks of these two values gives a score called the ASAP score, which is used to select the most likely partition. It is worth mentioning that OTUs delimitations obtained for two localities (Nouragues and Mitaraka) were confronted to morphological characters, and that in all cases the latter supported molecular delimitations (Decaëns et al. 2016, 2024), providing a strong argument for considering OTUs as relevant species proxies in further diversity analyses.

### Dataset and levels of sampling hierarchy

Data were organised into a community table containing the numbers of individuals collected for each OTUs (in columns) in each sample (i.e. each microhabitat sampled in each plot; in rows). This table was first used to produce a hierarchical partition of diversity according to a hierarchy of spatial extents or sampling levels (Table 1): 1) at the microhabitat scale, we considered the earthworms sampled in the different microhabitats in a given plot (i.e. soil, decomposing trunks, litter accumulations, streamside sediments and epiphytes); 2) at the community scale, we grouped the specimens collected from the various microhabitats within a given plot; 3) the habitat scale refers to specimens collected in different plots of the same habitat type within a given locality (i.e. plateau, slope and lowland forest, and inselberg habitats including hilltop forest, transitional forest and rocky savannah); 4) the landscape scale, groups together the data from all the plots sampled in the various habitats of the same locality; 5) the regional scale, corresponds to the spatial extent covered by the entire dataset.

**Table 1.**
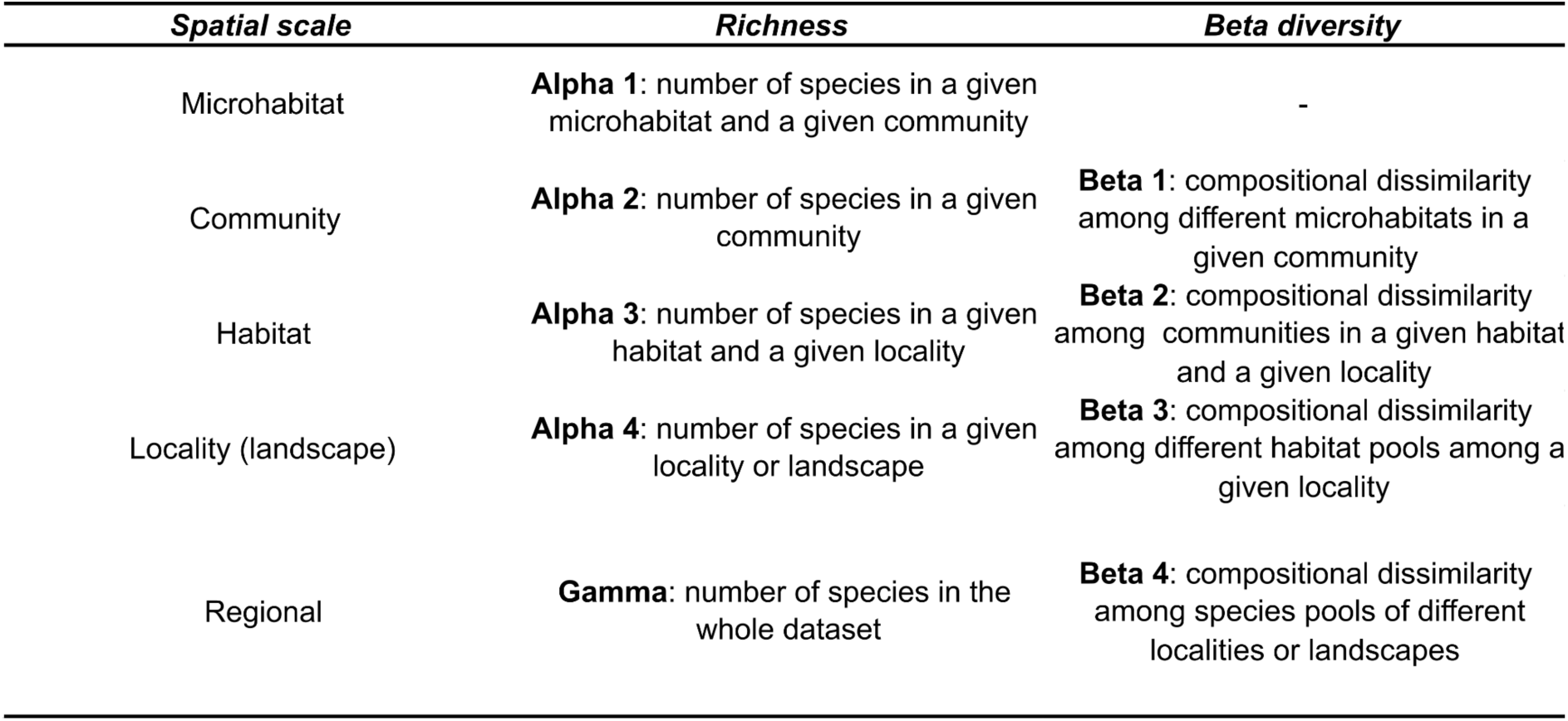
Different levels of sampling hierarchy considered and definition of the diversity measures considered in the calculations of richness and beta diversity in the diversity partitioning.

| *Spatial scale* | *Richness* | *Beta diversity* |
| --- | --- | --- |
| Microhabitat | **Alpha 1:** number of species in a given microhabitat and a given community | - |
| Community | **Alpha 2:** number of species in a given community | **Beta 1:** compositional dissimilarity among different microhabitats in a given community |
| Habitat | **Alpha 3:** number of species in a given habitat and a given locality | **Beta 2:** compositional dissimilarity among communities in a given habitat and a given locality |
| Locality (landscape) | **Alpha 4:** number of species in a given locality or landscape | **Beta 3:** compositional dissimilarity among different habitat pools among a given locality |
| Regional | **Gamma:** number of species in the whole dataset | **Beta 4:** compositional dissimilarity among species pools of different localities or landscapes |

### Multiscale diversity partitioning

The hierarchical partitioning of diversity aims to separate the relative contributions of the alpha and beta components to overall diversity (gamma) (Crist et al. 2003). Beta diversity can then be divided into hierarchical levels (i.e. spatial scales). To assess the robustness of our conclusions, we analysed our dataset using two complementary approaches: 1) an additive partitioning developed from the niche theory (Levins 1968, MacArthur 1968) and adapted statistically by Crist et al. (2003), and 2) a multiplicative partitioning using the approach proposed by Whittaker (1972). Both approaches share the objective of decomposing regional diversity, noted gamma (γ), into an inter-group component noted beta (β) and an intra-group component noted alpha (α), but differ fundamentally in the way they define beta diversity. For additive partitioning, β = γ – α (Wagner et al. 2000), whereas for multiplicative partitioning, β = γ / α (Whittaker 1972). Both partitioning are operational method for analysing species diversity across multiple spatial scales valid when restricted to species richness (Beck et al. 2012), as is the case in our work, even though they are not uncontested when applied to other metrics of diversity (Jost 2007, Baselga 2010, Marcon et al. 2014).

For both approaches, richness (i.e. ‘inventory diversity’ sensu Whittaker et al. 2001) is broken down as follows: alpha 1 corresponds to the number of OTUs observed at the microhabitat scale, alpha 2 at the community scale, alpha 3 at the habitat scale, and alpha 4 at the landscape scale. Regional or gamma diversity corresponds to the number of OTUs present in the entire dataset (Table 1).

For beta diversity, the partition is expressed as follows: beta 1 corresponds to inter-microhabitat or intra-community beta diversity, i.e. the compositional dissimilarity between the different microhabitats of a given community, beta 2 corresponds to intra-habitat beta diversity, i.e. between communities belonging to the same habitat in a given landscape, beta 3 corresponds to inter-habitat or intra-landscape beta diversity, i.e. between different pools of habitats in the same landscape, and beta 4 corresponds to the compositional dissimilarity between the different landscapes sampled over French Guiana (Table 1).

In the case of additive partitioning, the calculation of beta diversity at several sampling levels is as follows:

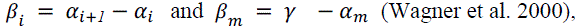

with partition levels varying from *i* to *m*, and *m* corresponding to the highest level before the regional scale.

Each value of additive beta diversity is then expressed as a percentage of gamma diversity to reflect the relative contribution of the different hierarchical levels to regional diversity.

In the case of multiplicative partitioning, the calculation of beta diversity at several sampling levels is as follows:

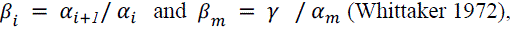

with partition levels varying from *i* to *m*, and *m* corresponding to the highest level before the regional scale.

The multiplicative beta diversity of a given level is maximum when all the surveys of that level contain entirely different species, and in this case it is equivalent to the number of surveys (Jost 2006). The values can then be standardised by expressing them as a fraction of maximum beta diversity (Glowacki et al. 2011), i.e. by dividing them by the actual number of situations corresponding to each sampling level.

Alpha and beta diversity values were compared with the results of a null model simulating, for each spatial scale, the values expected for assemblages made up of the same individuals as those found in the samples, but assembled by chance alone. The model used is a non-sequential algorithm for count matrices, which preserves the sum of the columns (i.e. the species frequency) and randomises the individuals between the cells in each column of the matrix. This type of null model is recommended for the analysis of species lists obtained from standardised samples taken from relatively homogeneous habitats (Gotelli 2000), which corresponds well to our dataset. More specifically, we performed 999 randomisations using the *’c0_ind’* algorithm of the *’commsim’* function in the *’vegan’* package for R-4.3.1 (Oksanen et al. 2019, R Core Team 2023). The diversity partitioning and the comparison between simulated and observed values were carried out using the *’adipart’* and *’multipart’* functions from the same package. For each scale, the function returns a standardised effect size (SES) of the observed statistic, with an associated p-value.

### Regional diversity estimation

To estimate the total number of species likely to exist in French Guiana, we divided the study region into 100 x 100-km^2^ cells in order to obtain as many contiguous cells with occurrence data as possible (i.e. six cells; SI Fig. S1). We organised the data into a contingency table showing the occurrences (i.e. presence/absence) of OTUs in each of these cells. This table was used to represent a rarefaction/extrapolation curve showing how OTUs accumulate as a function of the number of cells considered (intrapolation) and extrapolating the shape of the curve to a hypothetical dataset where the sampling effort would have been multiplied by two (extrapolation). We then calculated the asymptotic Chao 2 index, which can be used to obtain an approximation of the real number of species, as illustrated by a recent study on the global number of tree species (Cazzolla-Gatti et al. 2022). These analyses were performed using the *’iNEXT’* package for R (Hsieh et al. 2019); confidence intervals around the rarefaction/extrapolation curves and the Chao index were obtained using a bootstrap of 200 replications.

## Results

### Overall results

A total of 4729 earthworm specimens were collected in the 125 sampling plots and we obtained DNA barcodes for 3555 of them (i.e. 75.2%). The ASAP method enabled us to group these sequences into 256 OTUs, of which 25 were observed in Galbao, 34 in Itoupé, 27 in Kaw, 21 in Laussat, 34 in Mitaraka, 41 in Nouragues-Inselberg, 41 in Nouragues-Parare, 35 in Paracou, 44 in Saül and 30 in Trinité. Of these OTUs, 117 (i.e. 45.7% of the total) were represented only by juveniles or body fragments, and could not have been included in our study if identifications had been based solely on morphology. The most frequent OTU was represented by 476 individuals in the dataset, while 21.5% of OTUs corresponded to singletons (i.e. represented by a single individual) and the majority of OTUs (64.3%) were considered as rare since fewer than eight specimens have been collected (Fig. 2A). Similarly, 85.9% of OTUs were only sampled in a single locality (Fig. 2B) and only two OTUs (i.e. OTU#026 which corresponds to *Pontoscolex corethrurus* (Müller, 1857) and OTU#028 which corresponds to *Nouraguesia parare* Csuzdi & Pavlíček, 2011) were observed in the 10 study localities.

**Figure 2.**
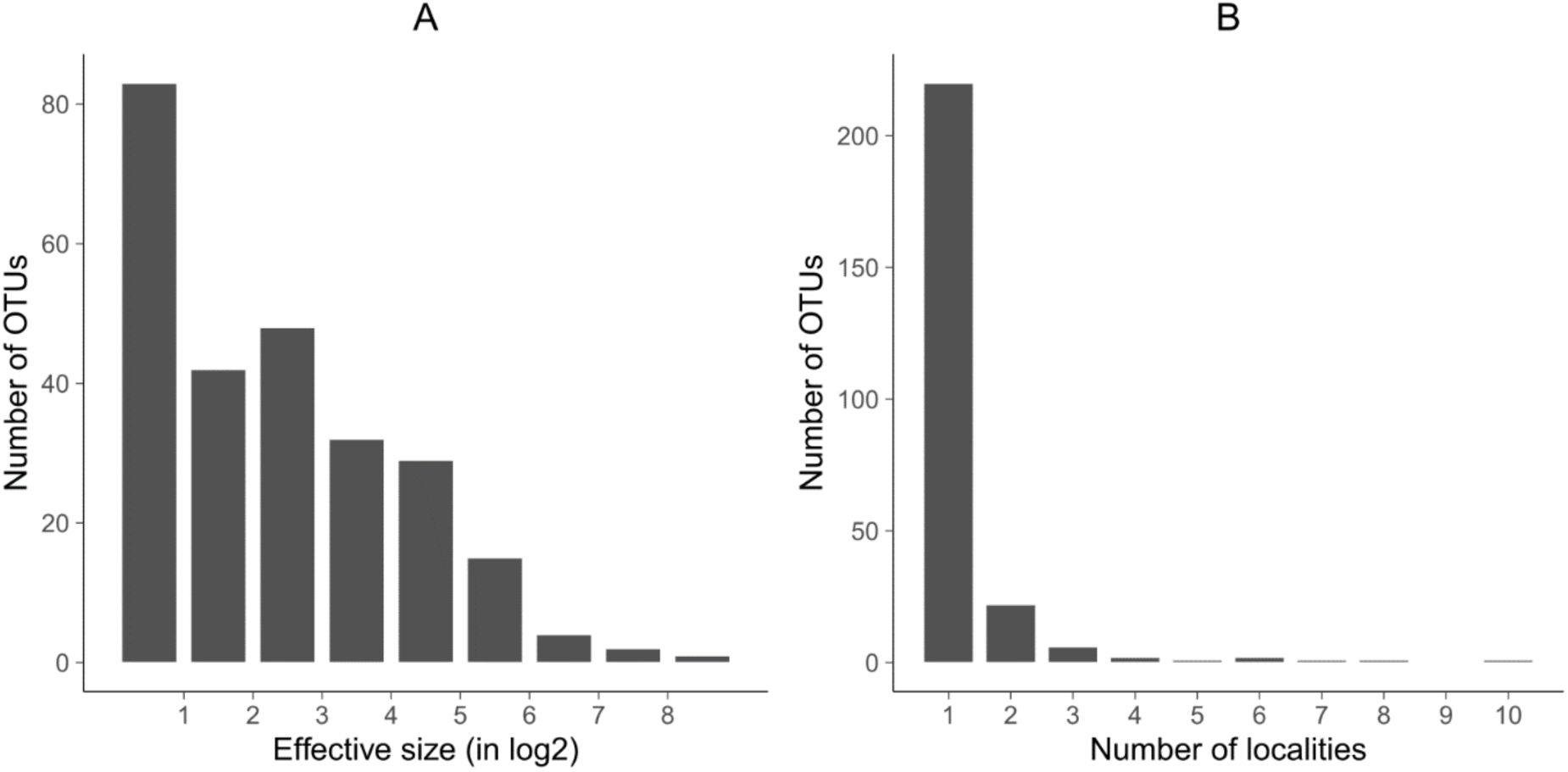
Distribution of earthworm abundance and occurrence in rainforests of French Guiana: A) Frequency histogram (Preston diagram) showing the number of OTUs observed per size class (in Log2); B) Occurrence histogram showing the number of OTUs per occurrence class (i.e. number of localities in which each OTU was observed).

### Hierarchical diversity partitioning

Richness averaged 3.9 OTUs at the microhabitat scale (alpha 1), 7 OTUs at the community scale (alpha 2), 15.9 OTUs at the habitat scale (alpha 3) and 33.2 OTUs at the landscape scale (alpha 4) (Fig. 3A). Generally speaking, the richness of the first two levels (alpha 1 and alpha 2) contributed only marginally to regional richness (e.g. 2.6% for alpha 2). For all scales, observed richness was significantly lower than expected from the null model, and the absolute value of the standardised effect size decreased steadily from alpha 1 to alpha 4 (Fig. 3B).

**Figure 3.**
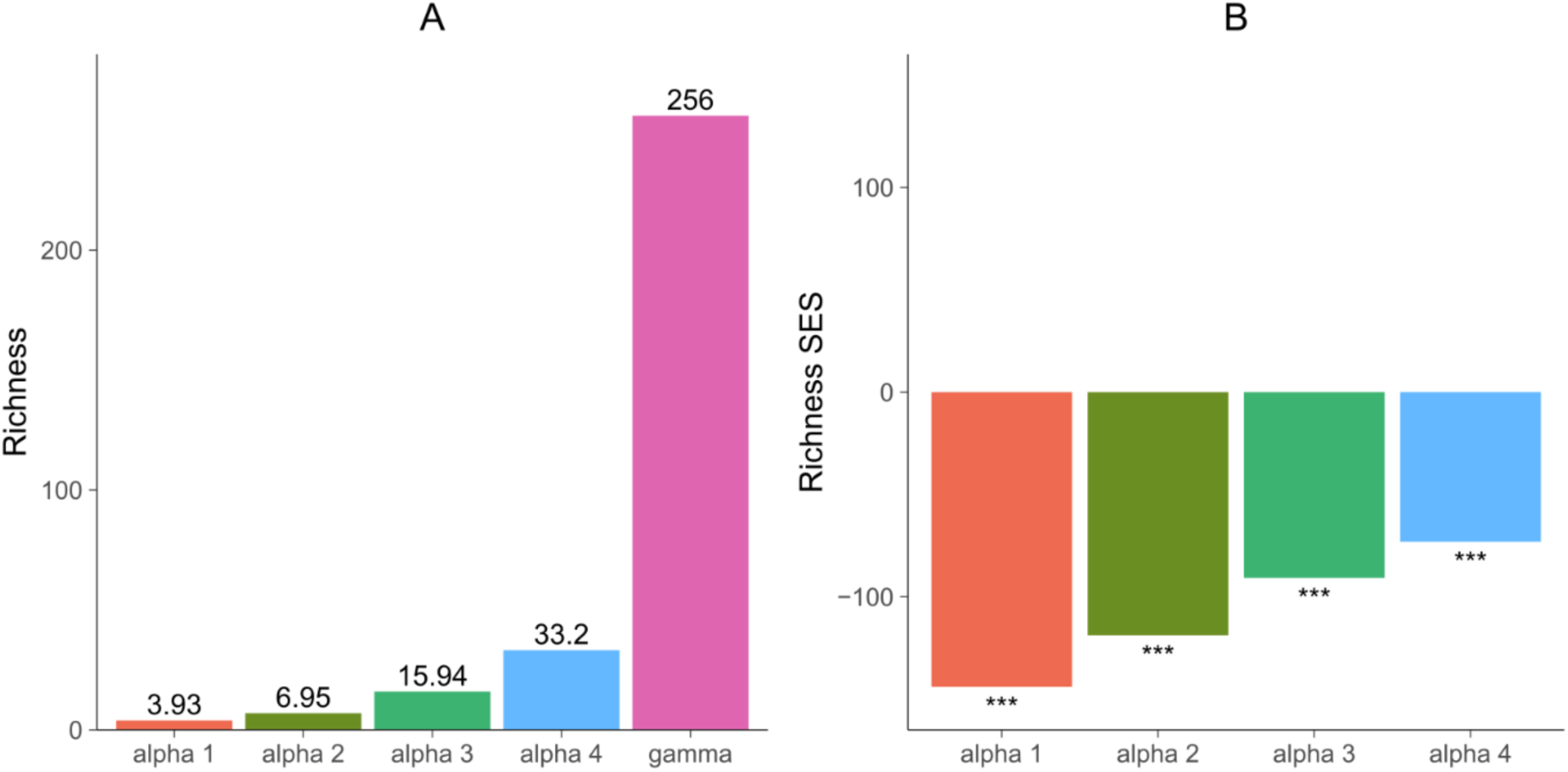
Earthworm OTU richness at different spatial scales in rainforests of French Guiana: A) Mean values observed at the microhabitat (alpha 1), community (alpha 2), habitat (alpha 3) and landscape (alpha 4) scales, and total number of species observed in the entire dataset (gamma); B) Standardised effect sizes (SES) associated with each of the alpha diversity measures. Chi-square significance codes: 0 ‘***’ 0.001 ‘**’ 0.01 ‘*’ 0.05 ‘.’ 0.01.

The additive diversity partitioning largely mirrors richness patterns, highlighting that 87% of regional diversity was explained by beta diversity between remote landscapes (beta 4, Fig. 4A). The relative importance of other hierarchical sampling levels was comparatively low, with compositional differences between microhabitats (beta 1), communities (beta 2) and habitats (beta 3) explaining only 1.2%, 3.5% and 6.7% of regional diversity respectively (Fig. 4A). This trend was confirmed by the results of the null model, since the values obtained for beta 1 to beta 3 were significantly lower than expected by chance, while the opposite was observed for beta 4 (Fig. 4B). When the multiplicative approach was used, beta diversity between microhabitats was the highest (beta 1 = 87.4% of maximum beta diversity), followed by beta diversity among distant landscapes (beta 4 = 77.1%), with the other two partition levels ranging from 59.6 to 64.9% (Fig. 4C). However, the values obtained were significantly lower than expected by chance for beta 1 and beta 2, while they were significantly higher than expected by chance for beta 3 and especially beta 4 (Fig. 4D).

**Figure 4.**
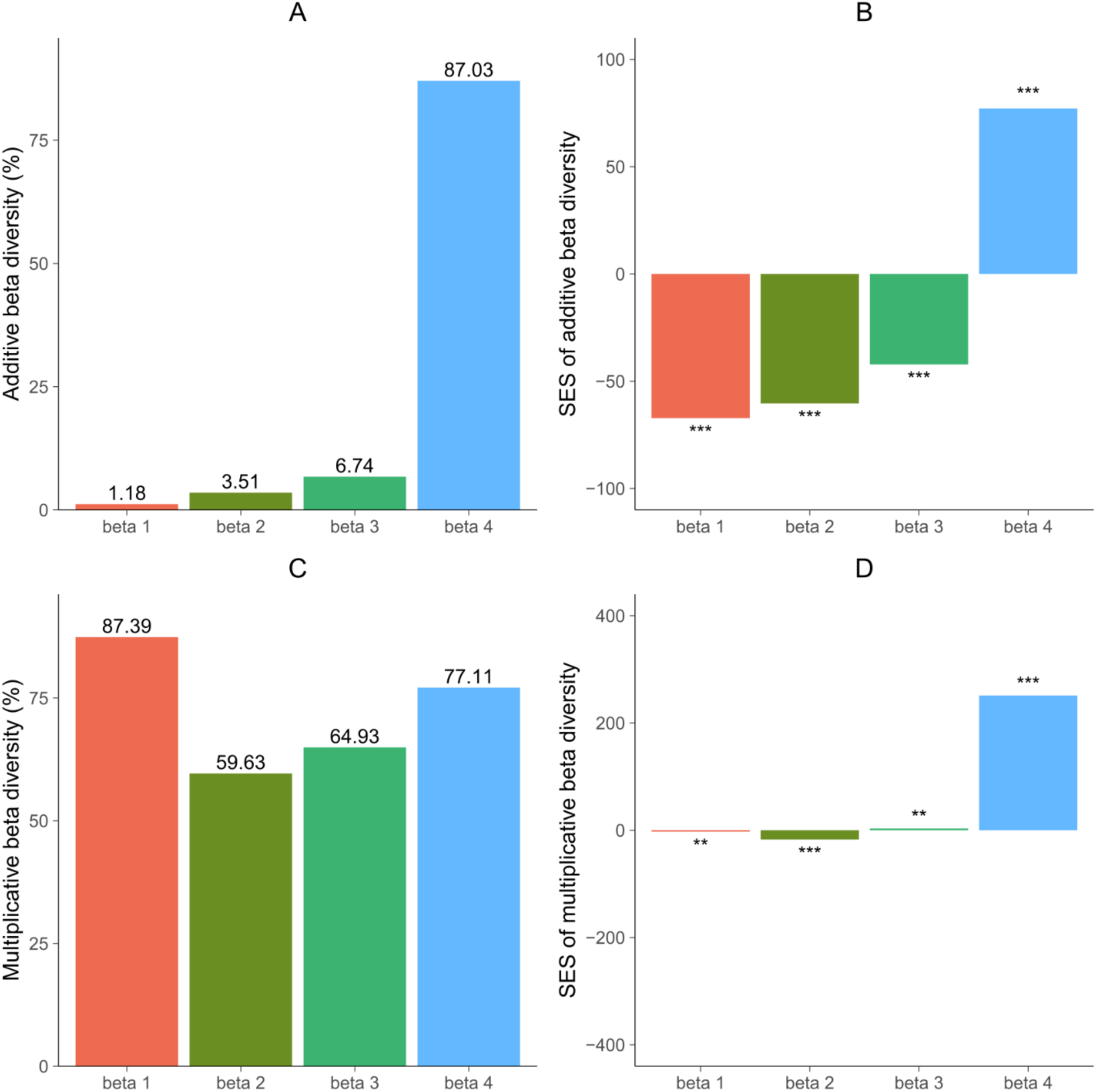
Additive and multiplicative partitionings of beta diversity of earthworm OTUs at different spatial scales in rainforests of French Guiana: A) proportion of gamma diversity (%) explained by additive beta diversity between microhabitats (beta 1), between communities of a given habitat (beta 2), between habitats (beta 3) and between landscapes (beta 4); B) Standardised effect sizes (SES) of additive beta diversity; C) Multiplicative partitioning of beta diversity (in % of maximum multiplicative beta diversity) for the same hierarchical levels as in A; D) Standardised effect sizes of multiplicative beta diversity. Chi-square significance codes: 0 ‘***’ 0.001 ‘**’ 0.01 ‘*’ 0.05 ‘.’ 0.01.

### Estimation of gamma diversity

The rarefaction curve shows that the completion of sampling on the scale of French Guiana was far from being achieved (Fig. 5). The asymptotic plateau on the curve was not even reached by extrapolating to a hypothetical dataset in which sampling effort would have been doubled, which suggests the existence of a significant species diversity that has not yet been sampled on a regional scale. This observation is confirmed by the calculation of the asymptotic Chao 2 index, which conservatively predicts that at least 1,725 species of earthworm could exist in total in French Guiana. The standard deviation associated with this index is high (i.e. +/− 280, i.e. 16.2% of the estimate), indicating that sampling effort is still too low to obtain a robust estimate, and that the figure obtained should be considered as a minimum value of the true diversity.

**Figure 5.**
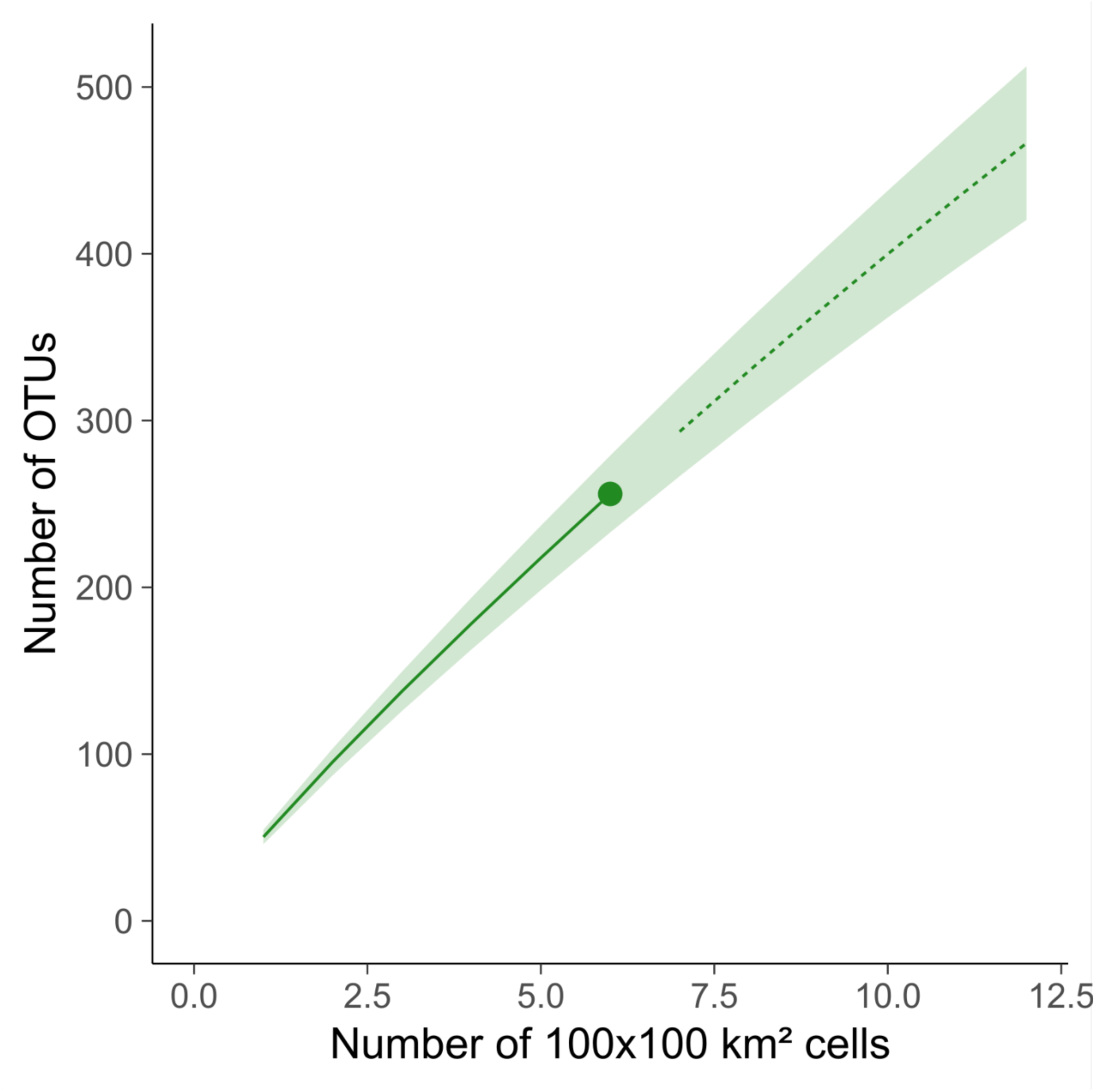
Rarefaction and extrapolation curve showing how observed OTUs accumulate as a function of the number of 100 x 100 km^2^ cells sampled. The solid line represents the rarefaction curve, the dotted line represents the extrapolation curve; the shaded area represents the 95% confidence interval based on a bootstrap with 200 replications.

## Discussion

### Beta diversity contribution to regional diversity

The hierarchical diversity partitioning supports our first hypothesis by clearly highlighting the importance of higher-order beta diversity in explaining the regional diversity of earthworms in French Guiana. The additive beta component at this scale (beta 4) explains 87% of the number of species observed regionally, which is much higher than expected by chance, while the contribution of lower hierarchical levels amounts to around 11.4%, and is lower than expected from the null model. This pattern is broadly confirmed by the multiplicative approach. Generally speaking, the importance of the beta component has been shown in a large number of studies involving a partitioning of tropical biodiversity, often with an increasing contribution along the hierarchy of scales considered. For example, it has been shown that regional diversity is essentially explained by beta diversity between biogeographical regions and between landscapes in moths in Southeast Asia (Beck et al. 2012) and in ants in Brazil (Marques and Schoereder 2014). Other studies have highlighted the importance of beta diversity between or within habitats, such as in spiders in Colombia (Cabra-Garcia et al. 2010) or ants in the Atlantic forests of Brazil (Sampaio et al. 2023), but they hardly compare with our work due to their more restricted geographical scope. Although they have not formally used a diversity partition, several studies have also demonstrated the importance of the beta component between regions and between habitats to explain the regional diversity of earthworms in Amazonia (Lavelle and Lapied 2003, Maggia et al. 2021, Conrado et al. 2023). Our study goes further by demonstrating unequivocally that the contribution of the beta component between different landscapes, rather than between habitats, sites or microhabitats, is by far the most important.

This result highlights the importance of regional mechanisms in shaping the spatial distribution of earthworm species diversity. This can be explained by the low dispersal capacity of these organisms, which may have favoured allopatric speciation events wherever small biogeographical barriers may have prevented earthworm movements (Lavelle and Lapied 2003). The isolation of populations and the resulting reduction in gene flow increases genetic differentiation between populations and the probability of local radiation events, as has been demonstrated in Amazonia for various animal and plant taxa (Boubli et al. 2015, Dambros et al. 2020, Fouquet et al. 2022). In French Guiana, it is likely that the dense hydrographic network may have played this role of vicariant agent, since the rivers are wide enough not to allow tree trunks to form natural bridges when they fall across the channel. Consequently, each interfluve delimited by rivers of sufficient size is likely to have created conditions favourable to the appearance of a specific pool of species, and most species have a restricted distribution range (Maggia et al. 2021).

Compared with the beta component between landscapes, lower hierarchical levels contribute significantly less to regional diversity. In particular, beta diversity between microhabitats (beta 1) is lower than expected by chance, whether calculated additively or multiplicatively, suggesting that earthworm communities tend to be made up of generalist species in terms of their use of microhabitats. This can be explained by the preponderance in the assemblages of epigeic pigmented species (Decaëns et al. 2016), which are among the most mobile and can be found both in the superficial soil layers and in above-ground microhabitats. Conversely, the results obtained for level 2 and 3 beta components suggest a habitat signal in community composition. In a given landscape, communities in the same habitats are more similar than expected by chance (negative beta 2 SES). On the other hand, although the additive contribution to regional diversity of the inter-habitat beta component (beta 3) remains low, beta diversity at this scale is nevertheless high compared with its theoretical maximum when calculated multiplicatively. This indicates a significant level of habitat pool specialisation, which is in line with the conclusions of other studies that have shown such a habitat signal within Amazonian earthworm communities (Decaëns et al. 2016, Maggia et al. 2021, Conrado et al. 2023).

### How many species in French Guiana?

The importance of beta components between distant landscapes and the shape of the rarefaction/extrapolation curves calculated from the occurrences of species in 100 x 100-km^2^ cells suggest a severe under-sampling at the regional scale, and raise the question of the number of species that could exist in French Guiana. The Chao 2 estimator predicts that at least 1,725 OTUs or species could be present in the study region. This estimate is a spectacular increase compared to the 33 species previously cited by Brown and Fragoso (2007), even though these authors already admitted that this number could not be representative of the real regional diversity. It is also much higher than the 250 species suggested by (Maggia et al. 2021), which was based on a smaller dataset than ours. The inflation of the estimate with the increase in sampling effort shows that the data are probably still insufficient to obtain an accurate estimate. Consequently, the calculation of the Chao index is still closely correlated with the size of the dataset, and any additional sampling effort in future work will probably result in an upward estimate of regional diversity.

This unambiguously illustrates the taxonomic deficit that characterises earthworms in French Guiana and, more generally, in tropical soils. A small proportion of the 256 OTUs sampled in our work have been formally described to date (Decaëns et al. 2024), and they represent only a small fraction of the regional diversity of French Guiana. An estimate of almost 1,725 species in an area as small as French Guiana also calls into question the commonly accepted estimate of 10,000 to 20,000 earthworm species on a global scale (Reynolds 1994, Decaëns et al. 2006, Blakemore 2009, Anthony et al. 2023). French Guiana represents just 1.5% of the surface area of the Amazonian rainforest, and a negligible fraction of the tropical rainforests as a whole. It will be interesting in the future to figure out whether the patterns of diversity highlighted in our study are repeated in other tropical regions. If this is the case, it would be a strong argument for considering earthworms as a hyper-diverse group that has not yet been recognised as such due to the taxonomic deficit.

## Conclusion

Our study provides new insights into the factors controlling the diversity of an essential component of soil biodiversity in a region that is also essential from a biodiversity perspective. In accordance with our first hypothesis, we observed high regional diversity in these organisms with low mobility, essentially explained by geographical turnover between species pools from one landscape to another. Regional factors linked to evolutionary processes and biogeographical history therefore seem to be the main candidate factors responsible for the hotspot of species diversity highlighted by our study. In comparison, regional diversity is only marginally attributable to the integration of samples from contrasted habitats with very different species communities. Our results also support our fourth hypothesis, according to which French Guiana is home to a large number of as yet undiscovered species, and that this unsuspected regional diversity is likely to challenge the current consensus on the number of species existing on a global scale.

In addition to these advances in our fundamental understanding of how earthworm diversity is structured, our study argues for these organisms to be given greater consideration in tropical biodiversity management strategies. With nearly 2,000 species, earthworms would constitute a group equivalent in species number to terrestrial vertebrates (with around 1500 sp in French Guiana), and only surpassed by plants (with 5500 sp), beetles and lepidoptera (each with more than 5500 sp) (Combou 2011, Brulé and Touroult 2014). The unique distribution of this diversity, and in particular the remarkable importance of geographical turnover between distant localities, deserves to be considered in the definition of strategies for the protection of natural areas on the scale of French Guiana. Furthermore, the taxonomic deficit highlighted in our work is considerable, and the rate of description of new species could be much higher than for most hyper-diverse insect groups (Brulé and Touroult 2014), if we are able to maintain a substantial taxonomic effort in the future. This observation alone provides justification for mobilising the necessary resources to step up the sampling effort and enable newly discovered species to be described.

## Acknowledgements

Part of the dataset used in this study was acquired as part of the DIADEMA (Dissecting Amazonian Diversity by Improving a Multiple Taxonomic Group Approach), DIAMOND (Dissecting and Monitoring Amazonian Diversity) and Wormbank (DNA barcoding earthworms in biodiversity hot spots of French Guiana) projects funded by “Investissement d’Avenir” grants managed by the Agence Nationale de la Recherche (CEBA: ANR-10-LABX-25-01; TULIP: ANR-10-LABX-41). Sampling in Mitaraka was carried out as part of the “Our Planet Reviewed” French Guiana 2015 expedition (Touroult et al. 2018) organised by the Muséum national d’Histoire naturelle (MNHN, Paris) and Pro-Natura international in collaboration with the Amazonian Park of French Guiana, and financed by the European Fund for Regional Development (FEDER), the Regional Council of French Guiana, the General Council of French Guiana, the Direction de l’Environnement, de l’Aménagement et du Logement and by the Ministère de l’Éducation nationale, de l’Enseignement supérieur et de la Recherche. At the Réserve naturelle des Nouragues, the project was supported by two Centre National de la Recherche Scientifique (CNRS) Nouragues grants in 2010 and 2011. At the Réserve naturelle de la Trinité, part of the funding was provided by the nature reserve. The authors would like to thank the Parc Amazonien de Guyane (http://www.parcamazonien-guyane.fr), the Réserve naturelle de la Trinité (http://www.reserve-trinite.fr/) and the Réserve naturelle des Nouragues (http://www.nouragues.fr/) for authorising access and collecting. Sample DNA barcoding received funding from the Canadian Centre for DNA Barcoding (CFREF-2015-00004), as well as from the CNRS through the BC-Wormbank project (APEGE 2013 call).

## Author Contributions

Conceptualisation, AG, MH, TD; methodology, AG, EM, MEM, MH, PG, TD; field work– data acquisition: EL, TD; formal analyses, AG, MEM, TD; writing–original draft, AG; writing–review and editing, EL, EM, MEM, MH, PG, TD; supervision: MH, TD; funding acquisition: TD. All authors have read and agreed to the published version of the manuscript.

## Conflict of interest statement

The authors declare no conflicts of interest.

## Data availability statement

The complete list of specimens, associated metadata, COI sequences and GenBank accession numbers are available in the BOLD dataset “Earthworms from French Guiana – 2023 update” (DS-EWFG2023; doi.org/10.5883/DS-EWFG2023). The complete list of specimens and the community table are also deposited on the Zenodo repository (doi.org/10.5281/zenodo.10908657).

## Notes

https://doi.org/10.5883/DS-EWFG2023

https://doi.org/10.5281/zenodo.10908657

### Competing Interest Statement

The authors have declared no competing interest.

