## Appendix S1. Materials and methods supporting information for "Dissecting earthworm diversity in tropical rainforests"

*Table S1.* List of sampling localities, average geographical position, sampling periods and number of one-hectare plots sampled in each.

| Localities | Latitude | Longitude | Sampling date | Number of sampling plot |
|---|---|---|---|---|
| Laussat | 5.48117 | -53.5762 | 02/2014 - 02/2018 - 12/2019 | 7 |
| Paracou | 5.25833 | -52.9346 | 06/2013 | 10 |
| Trinite | 4.61871 | -53.4081 | 04/2016 | 10 |
| Kaw | 4.54064 | -52.2152 | 06/2013 - 04/2016 | 9 |
| Inselberg | 4.08133 | -52.6731 | 01/2011 - 06/2011 - 02/2018 | 25 |
| Parare | 4.02832 | -52.6778 | 06/2011 - 07/2011 - 02/2018 | 12 |
| Galbao | 3.60403 | -53.268 | 01/2016 - 01/2019 | 11 |
| Saül | 3.5587 | -53.2222 | 10/2013 - 02/2014 | 12 |
| Itoupe | 3.02696 | -53.079 | 01/2016 | 11 |
| Mitaraka | 2.243 | -54.465 | 03/2015 | 18 |

*Table S2.* Number of one-hectare plots sampled per habitat in each locality.

| Localities | Lowland forest | Slope forest | Plateau forest | Inselberg habitat | White sands |
|---|---|---|---|---|---|
| Laussat | 2 | - | 2 | - | 3 |
| Paracou | 3 | 4 | 3 | - | - |
| Trinite | 3 | 2 | 3 | 2 | - |
| Kaw | - | 2 | 7 | - | - |
| Inselberg | 2 | 10 | 8 | 5 | - |
| Parare | 4 | 3 | 5 | - | - |
| Galbao | 2 | 4 | 2 | 3 | - |
| Saül | 4 | 4 | 4 | - | - |
| Itoupe | - | 4 | 7 | - | - |
| Mitaraka | 3 | 3 | 4 | 8 | - |

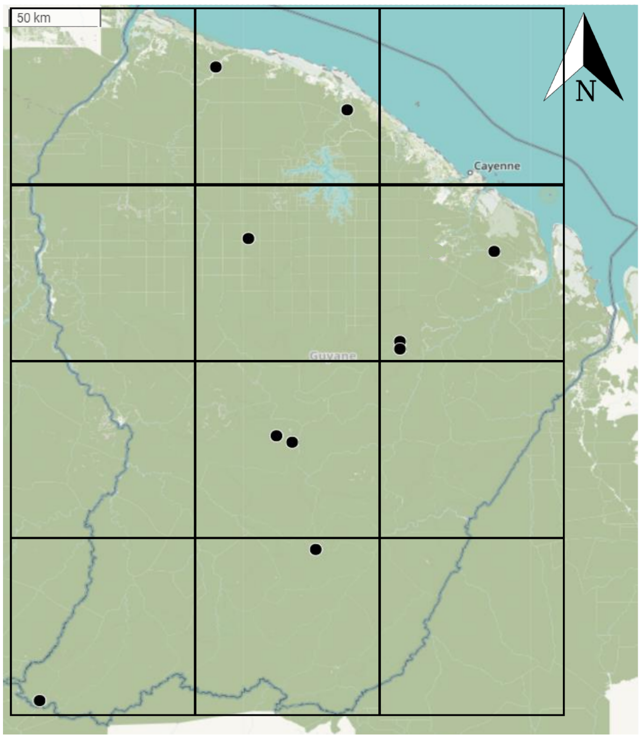


*Figure S1.* Division of French Guiana into 100 x 100 $km^2$ cells used to estimate diversity on a regional scale. Each point on the map represents a sampling locality as shown in Fig. 1.
